# Vitrifying multiple embryos in different arrangements does not alter the cooling rate^⋆^

**DOI:** 10.1101/2021.05.12.443816

**Authors:** Timothy Ostler, Thomas E. Woolley, Karl Swann, Andrew Thomson, Helen Priddle, Giles Palmer, Katerina Kaouri

## Abstract

Vitrification is the most common method of cryopreservation of gametes in fertility clinics due to its improved survival rates compared to slow freezing techniques. For the Open Cryotop^®^ vitrification device, the number of oocytes, or embryos, mounted onto a single device can vary.

In this work, a mathematical model is developed for the cooling of oocytes and embryos (samples). The model is solved computationally, to investigate whether varying the number of samples mounted onto the Open Cryotop^®^ affects the cooling rates, and consequently the survival rates, of vitrified samples. Several realistic spatial arrangements of samples are examined, determining their temperature over time. In this way we quantify the effect of spatial arrangement on the cooling rate.

Our results indicate that neither the spatial arrangement nor the number of mounted samples has a large effect on cooling rates, so long as the volume of the cryoprotectant remains minimal. The time taken for cooling is found to be on the order of half a second, or less, regardless of the spatial arrangement or number of mounted samples. Hence, rapid cooling can be achieved for any number or arrangement of samples, as long as device manufacturer guidelines are adhered to.

## 1. Introduction

Vitrification is ‘the supercooling of a liquid to a temperature at which the viscosity is so high that it can be defined as being at a solid state’ [4]. Vitrification has been used in the application of cryopreservation of gametes and embryos for decades [28] and is now the dominant method of cryopreservation in fertility, taking over from slow freezing methods [38]. High rates of cooling are considered important to the survival of vitrified oocytes and embryos [23, 31], as are high rates of warming during thawing after storage [34]. The process involves the application of permeating cryoprotectants in high concentrations, which mitigate the risk of formation of ice crystals [4, 35], and direct exposure to liquid nitrogen [18]. A number of different devices and techniques have been developed to facilitate this process, including open pulled straws [36], cryoloops [26], electron microscope grids [27] and the Open Cryotop^®^ device [23]. Vitrified oocytes or embryos are then transferred to storage until needed for an IVF cycle, or until some maximum time period has elapsed. The maximum period depends on national regulations; in the UK, this is governed by the HFEA, who currently suggest a maximum of 10 years [15].

Attempts have been made to study how the characteristics of various cryopreservation devices affect the rate of temperature change, often using the measure of ‘cooling rates’. The cooling rate is calculated as the change in temperature divided by the time taken to observe this change. Cooling rates are frequently reported for different temperature ranges, such as from 293.15K to 153.15K [25], 293.15K to 143.15K [31] and 253.15K to 173.15K [23]. Methods have been explored to measure cooling rates using thermocouple devices [22], and cooling rates are reported for several devices [36, 37]. An alternative approach to explore temperature evolution in such systems is through computational modelling, an established technique for vitrification devices [31, 32, 33]. Such thermal modelling is predominantly aimed at determining system parameters to fit data, or to give theoretical comparisons of different vitrification devices, in order to relate cooling and warming rates to the likelihood of embryo and oocyte survival.

In some cryopreservation techniques, multiple oocytes, or embryos (referred to as ‘samples’ hereafter, see Section 2.2.2), may be loaded onto a single device. For example, the Open Cryotop^®^ (Kitazato^®^) may have one to four samples [20], whilst the S-Cryolock^®^ device (Biotech Inc.) may carry one or two [17]. Recent literature has reported that post-thaw survival rates of embryos are affected by the number of embryos vitrified with a single cryo-carrier, but states that there is still further work required to determine some optimum number, and does not give an explanation for this effect [3]. Additionally, when multiple samples are loaded onto a single device, it has been observed that they can either remain isolated from one another, or arrange themselves in a more complex geometry. This variability in the spatial arrangements of samples leads to variability in practice between clinics, operators and even differences between procedures performed by the same operator. Thus, the question arises on whether such differences have any clinical significance, with respect to the survival of the samples. As far as the authors are aware, there is currently no research quantifying the influence of the number of plated samples and/or their spatial arrangement on the cooling rates of the vitrification process; this is what we set out to do in this work.

The objective of this study is to investigate whether the number of samples, in addition to their spatial arrangement on the vitrification device, affects cooling rates, through the application of computational modelling. In particular, the study focuses on the Open Cryotop^®^ device, with the vitrification protocol defined according to the user manual [20], although the approach taken in this study can be generalised for other similar devices.

## 2. Materials and Methods

### 2.1. Mathematical model of vitrification on the Open Cryotop^®^ device

The Open Cryotop^®^ device consists of a polypropylene plate with a depth of 0.1mm and a width of 0.7mm (see Figure 1) [19]. A sample, suspended in a cryoprotectant solution, is pipetted onto the end of the plate (Figure 2), and the entire construction is subsequently submerged into a liquid nitrogen bath.

**Figure 1:**
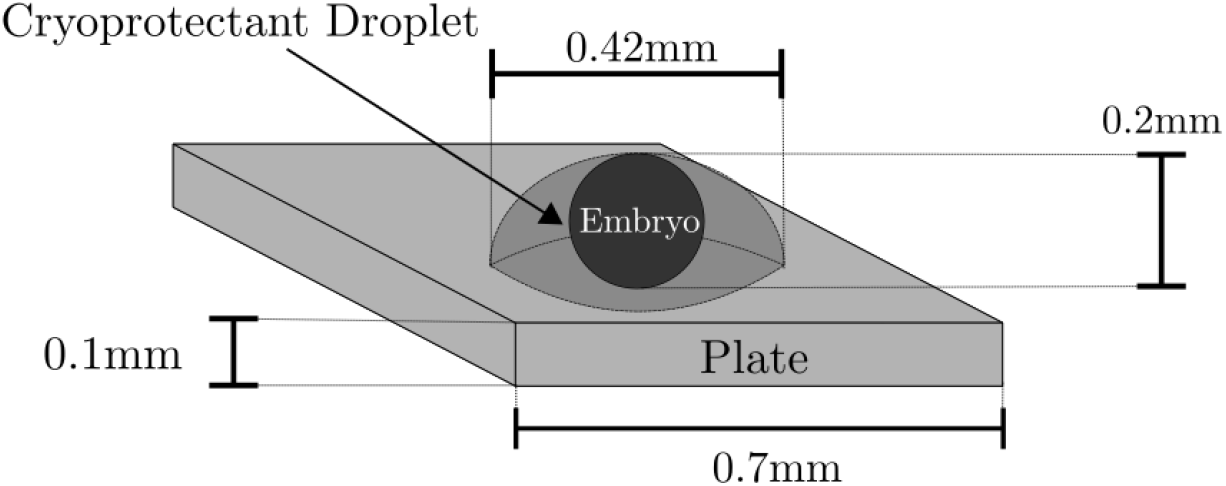
A schematic diagram of the thermal model, including dimensions of the three key domains: the sample, the plate and the cryoprotectant droplet. The entire system is submerged in liquid nitrogen.

**Figure 2:**
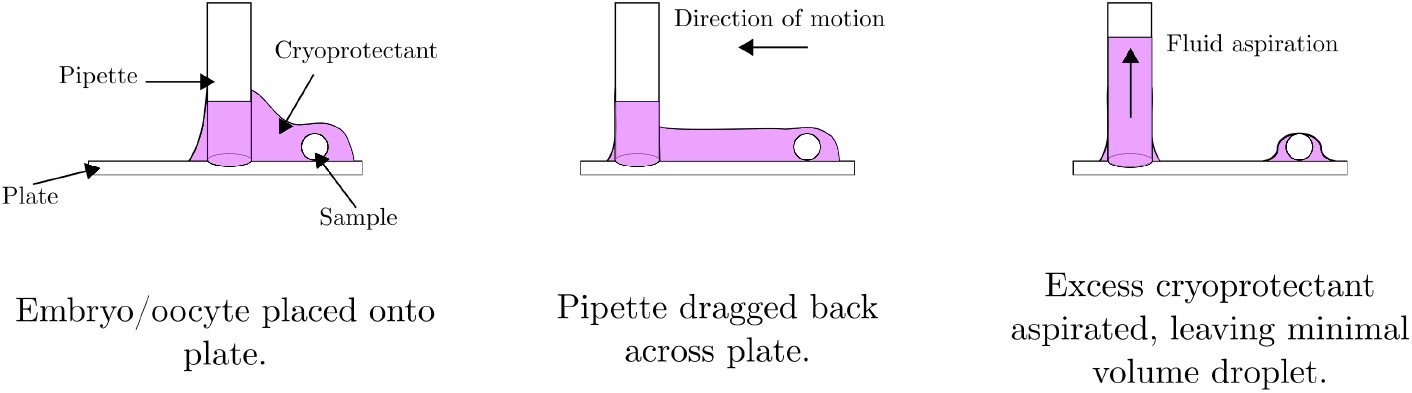
Diagram depicting the process of transferring a sample from a dish onto the plate of the Open Cryotop^®^, and removing excess cryoprotectant through aspiration to leave a ‘minimal droplet’. Adapted from the Open Cryotop^®^ user manual [20].

A schematic diagram of the system is presented in Figure 1. When modelling multiple samples, the samples are placed on the plate at specific locations. Then, cryoprotectant droplets are constructed around each sample, ignoring any overlap between droplets (see Figure 3, where an example with two droplets is constructed using COMSOL Multiphysics 5.5). Thus, the cumulative volume of cryoprotectant does not grow in proportion to the number of samples vitrified, as the overlap volume between droplets depends on how closely the samples are placed to one another and in which pattern. In this way, clinical practice is replicated in the modelling as excess cryoprotectant is aspirated, rather than transferring each sample in a fixed volume.

**Figure 3:**
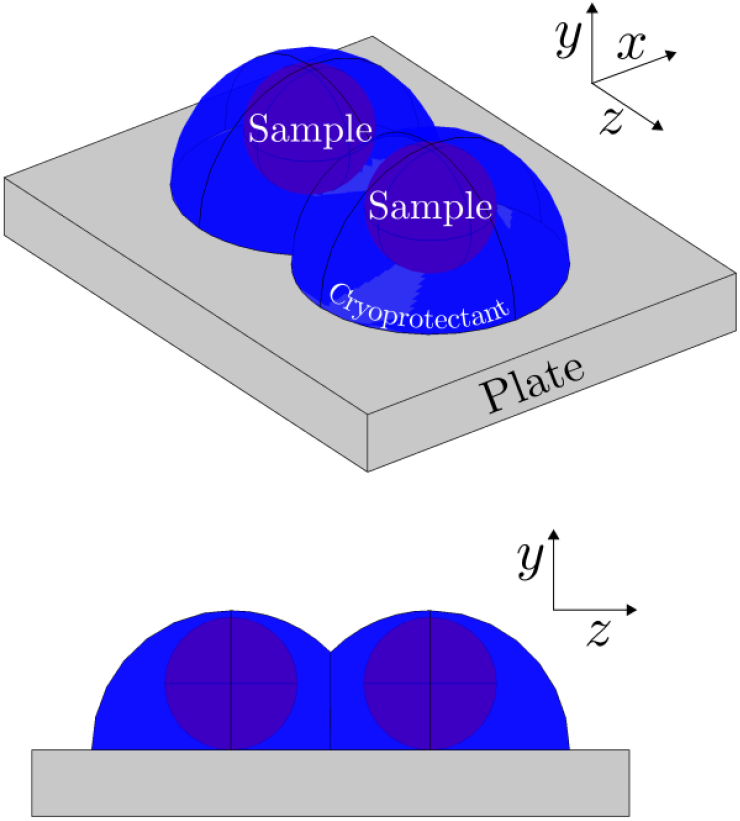
Two merged droplets rendered in COMSOL Multiphysics 5.5 droplets. The same set-up is displayed from two different angles.

Once the device, cryoprotectant droplet and sample are constructed, the temperature of each component is determined over time by solving the heat equation for the system. The heat equation is used [9], which states that the temperature *u* = *u*(*x, y, z, t*) of a component at some three dimensional location (*x, y, z*) and time *t* satisfies the partial differential equation,

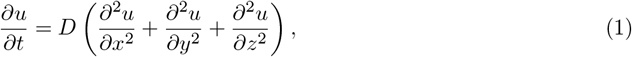

with an initial temperature of *u*(*x, y, z*, 0) = 310.15K. The diffusion coefficient, *D*(*u*) (*μ*m^2^/s), of the material is a positive scalar defined by the relation [9]

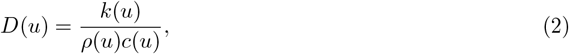

where:

- *c*(*u*) is the specific heat capacity of the material at temperature *u*, (J/(kg K)),
- *k*(*u*) is the thermal conductivity of the material at temperature *u*, (W/mK),
- *ρ*(*u*) is the density of the material at temperature *u*, (kg/m^3^).

Additionally, the rate at which liquid nitrogen cools the boundaries of the system may be described via a convective heat flux boundary condition [9],

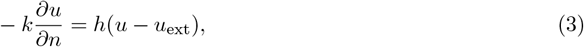

where *u*_ext_ is the temperature of the liquid nitrogen. The parameter *k* is the thermal conductivity of either the polypropylene plate or the sample in contact with the liquid nitrogen, and *h* (W/m^2^*K*) denotes the surface heat transfer coefficient, which defines the rate at which the sample, or the plate, transfers heat to the surrounding liquid nitrogen.

Whilst *D*(*u*) may be considered as a function of temperature [33], other approaches assume *k, ρ* and *c* (and thus *D*) are constant [31]. This assumption will be further addressed in Section 2.2.3.

The model incorporates the following domains (1):

- Liquid nitrogen, *Ln*,
- The cryoprotectant fluid, *F*,
- Samples, *S*, which is a disconnected set of spherical samples,
- The polypropylene plate, *P*.

Where two domains, *A* and *B*, share an interface, this interface is denoted by ∂Ω_*A,B*_. A cross section of the model is displayed in Figure 4 with labelled domains, interfaces and boundaries.

**Figure 4:**
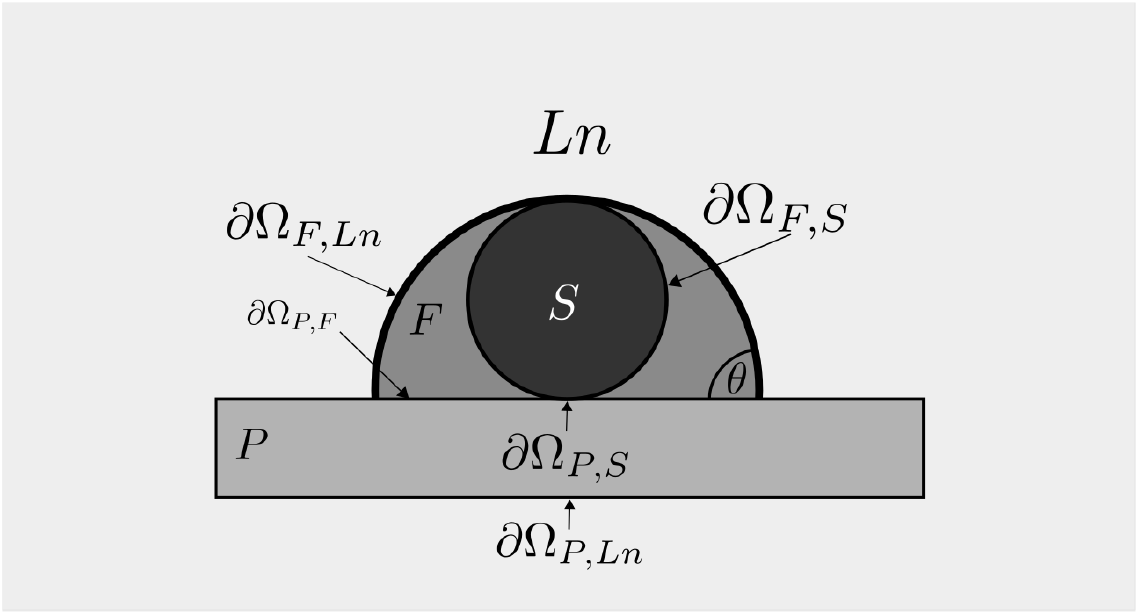
A cross section of the model geometry.

Let *u_P_* = *u_P_*(*x, y, z, t*), *u_F_* = *u_F_*(*x, y, z, t*) and *u_S_* = *u_S_*(*x, y, z, t*) be the temperatures in the plate, cryoprotectant fluid and the samples, respectively, at location (*x, y, z*) and time *t*. The temperature in the system is described by the following system of equations:

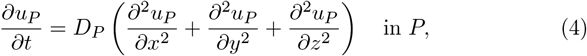

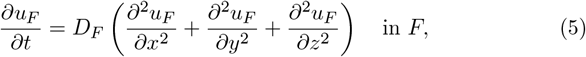

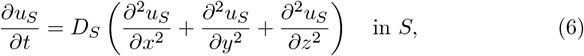

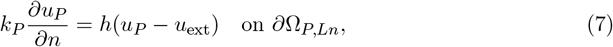

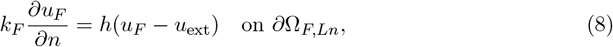

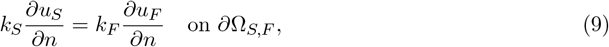

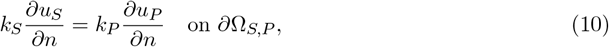

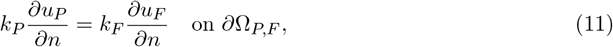

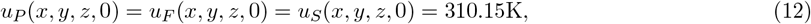

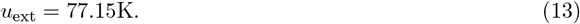

Equations (4)–(6) describe heat movement within individual domains. Equations (7) and (8) describe heat convection out of the system through the boundaries of the plate and cryoprotectant into the liquid nitrogen. Equations (9)–(11) describe conduction between the samples and cryoprotectant, samples and plate, and plate and cryoprotectant, respectively. Equations (12) and (13) describe the initial temperature and external temperature, respectively.

Cooling rates are calculated as the change in temperature divided by the time taken for that change to occur. We report cooling rates between an upper temperature of 310.15K, in accordance with our initial condition, and a lower limit of 143.15K, which is consistent with previous literature [31, 25]. The lower limit coincides with the glass transition temperature (*T_g_*) of the material, which is the temperature at which vitrification is considered to occur, as reported for similar cryoprotectant solutions [30].

### 2.2. Assumptions

Models enable us to represent reality with mathematical objects. Developing a mathematical model is a complex task that requires a number of simplifying assumptions. The modelling assumptions made in this study are discussed in the following sections to highlight their validity.

#### 2.2.1. Liquid nitrogen is a stationary, isothermal liquid

Liquid nitrogen is assumed to be a stationary, isothermal liquid with a fixed temperature of 77.15K. Hence, the fluid properties of the liquid nitrogen are not modelled in detail. This assumption is consistent with current modelling approaches [31], and is valid when the device is submerged far beneath the surface of the liquid nitrogen such that it is unaffected by bubble formation and bursting in the liquid nitrogen. Furthermore, the volume of the liquid nitrogen is much greater than the volume of the Open Cryotop^®^ device, thus energy transferred from the device to the liquid nitrogen is quickly spread throughout the system. Fluid motion would increase the rate of cooling, thus these results offer an upper estimate of the times taken for vitrification. Hence, our results offer conservative insights from a ‘worst case’ scenario.

#### 2.2.2. *Samples are spheres of radius* 0.1*mm*

Oocytes can vary in size and shape, but are typically spherical, with a diameter of approximately of 0.12mm [13]. Including the Zona Pellucida and perivitelline space, the total system diameter is approximately 0.15mm [39]. Cumulus cells are not included, as these are removed according to the Open Cryotop^®^ user manual [20]. Only uniform samples are considered, as deviation from a spherical shape is considered a form of dysmorphia [14]. The choice of 0.1mm is a conservative size estimate, and accounts for loss of water volume and artificial collapse during the vitrification process. Assuming a conservative size estimate for samples allows more samples to fit together in dense arrangements. Denser arrangements are more likely to give slower cooling rates, so this assumption also offers conservative insights in the time taken for vitrification.

#### 2.2.3. Thermophysical Properties and Parameters

Since the size of the Zona Pellucida does not change throughout the early stages of the development of an embryo, the same model may be used to describe oocytes and embryos. This makes the assumption that the structural changes occurring post fertilisation do not affect the thermal properties of the embryo, compared to the oocyte.

To construct a numerical model, parameter values must be chosen for *c, k* and *ρ* for the plate and cryoprotectant. These values are ‘characteristic of, and measured by, different experimental situations’ [9], and can only be accurate if measured for the specific case in question. The vitrification solution contains 15% ethylene glycol and dimethyl sulphoxide (DMSO) v/v each, as well as a 0.5M sugar solution [19]. The concentration of solutions of ethylene glycol in water has a documented effect on the thermal properties of the system [6]. Although experimental derivations of temperature dependent thermophysical properties exist for various compounds and tissues [10], parameters for the exact proprietary solution are not well documented, so it is necessary that some simplifying assumptions are made [31, 33]. Where temperature dependent data is available for similar compounds, it is possible to assume the cryoprotectant is made up of this compound, and interpolate between known recorded parameter values, as has been used in the separate problem of device warming [33]. Experimental recordings of chemical and tissue thermophysical properties are often incomplete, however, with only a few experimentally derived temperatures over a smaller temperature range than the cooling observed during vitrification [10]. The most complete dataset available is the temperature dependent thermophysical properties of water [5, 10, 24], so we undertake simulations using this set of parameters. Assuming the cryoprotectant is water is imperfect, as this requires that at 273.15K there is a phase change as the water freezes to ice, which the purpose of vitrification is to avoid, but this is the best data currently available in literature, and this assumption will act as a useful comparison for the effect of variable thermophysical parameters on the model.

In general, *k*(*u*) increases as temperature decreases, whilst *c*(*u*) and *ρ*(*u*) decrease [10], which by Equation (2) implies *D*(*u*) increases with falling temperature. This increase in *D*(*u*) is shown to cause overestimation of cooling times under constant parameter assumptions [10]. In the absence of suitable temperature dependent data, it can be necessary to assume that thermophysical parameters are constants [31], noting that as constant parameter assumptions underestimate cooling rates, any simulated arrangement of samples generating suitable cooling rates under the constant parameter assumption will also have sufficiently high cooling rates in the laboratory. In this work, we use two sets of constant parameters. The first set assumes the cryoprotectant to be comprised of ethylene glycol only, which uses some intuition that the vitrification solution should behave more similarly to the cryoprotectants it contains than water, or they would serve no purpose in the vitrification process. The solution is still water based, however, so as a comparison we assume a second set of parameters in which the cryoprotectant consists of vitreous water, which is consistent with previous literature approaches [31].

Constant values for *c, ρ* and *k* are reported in Table 1 for ethylene glycol and vitreous water, whilst variable thermophysical properties of water are shown in Table 2. During simulations, linear interpolations are used to generate parameters between recorded values, and outside of the reported temperature range, thermophysical parameters are extrapolated as constants. The diffusion coefficients for all three of the permutations of this assumption are displayed in Figure 5.

**Table 1:**
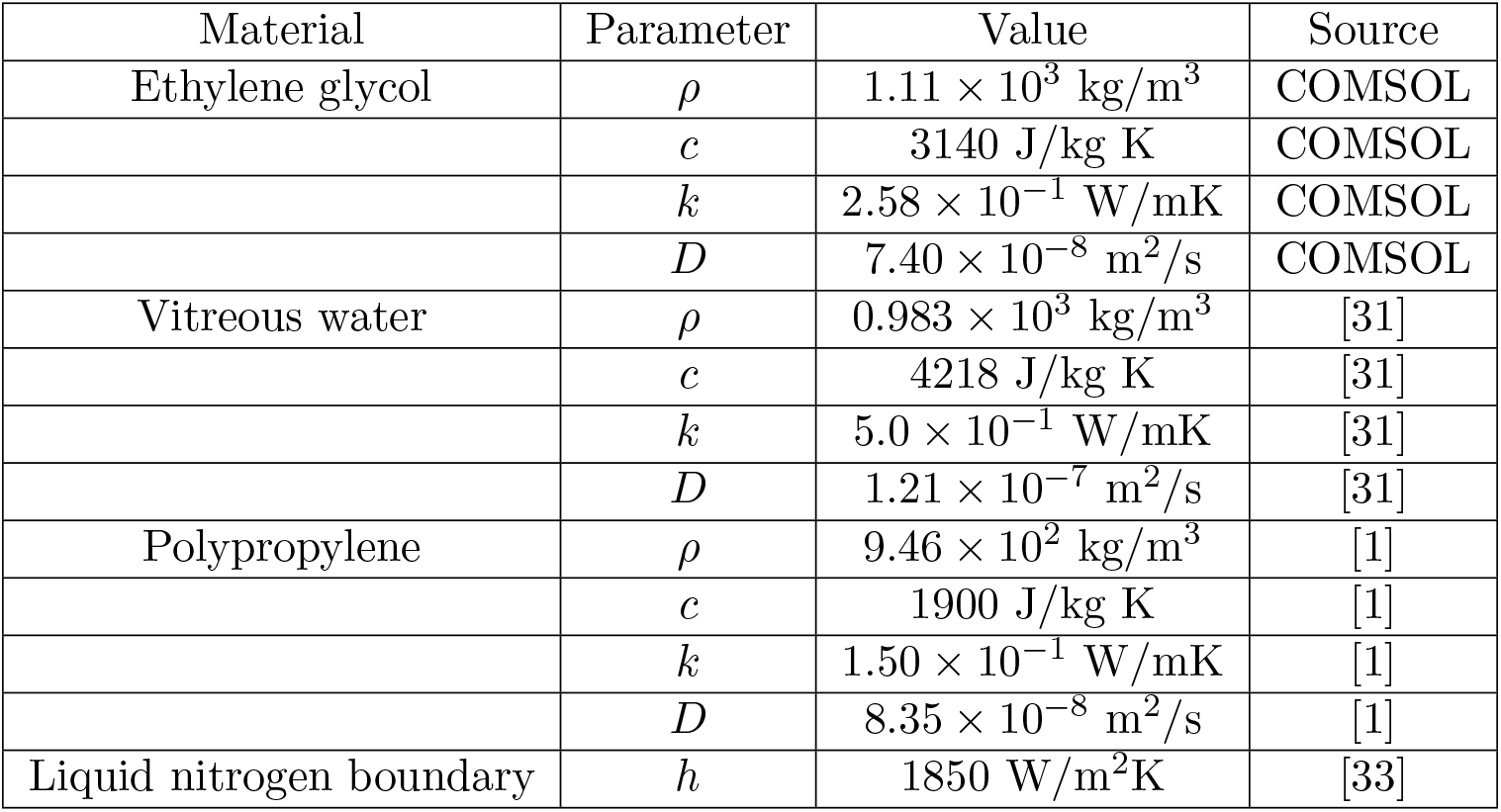
The thermal properties for the materials involved in the system.

| Material | Parameter | Value | Source |
| --- | --- | --- | --- |
| Ethylene glycol | $\rho$ | $1.11 \times 10^3 \text{ kg/m}^3$ | COMSOL |
| | $c$ | $3140 \text{ J/kg K}$ | COMSOL |
| | $k$ | $2.58 \times 10^{-1} \text{ W/mK}$ | COMSOL |
| | $D$ | $7.40 \times 10^{-8} \text{ m}^2/\text{s}$ | COMSOL |
| Vitreous water | $\rho$ | $0.983 \times 10^3 \text{ kg/m}^3$ | [31] |
| | $c$ | $4218 \text{ J/kg K}$ | [31] |
| | $k$ | $5.0 \times 10^{-1} \text{ W/mK}$ | [31] |
| | $D$ | $1.21 \times 10^{-7} \text{ m}^2/\text{s}$ | [31] |
| Polypropylene | $\rho$ | $9.46 \times 10^2 \text{ kg/m}^3$ | [1] |
| | $c$ | $1900 \text{ J/kg K}$ | [1] |
| | $k$ | $1.50 \times 10^{-1} \text{ W/mK}$ | [1] |
| | $D$ | $8.35 \times 10^{-8} \text{ m}^2/\text{s}$ | [1] |
| Liquid nitrogen boundary | $h$ | $1850 \text{ W/m}^2\text{K}$ | [33] |

**Table 2:** Thermophysical properties of water, as reported by [24]. Two values are reported for 273.15K depending on whether modelling ice or water, so to allow for linear interpolation, we report the values of ice at 273.14K and the values for water at 273.16K.

| Temperature | $k$(W/mK) | $\rho$ (kg/m<sup>3</sup>) | $c$ (J/kg) K | $D$ (m<sup>2</sup>/s) |
| --- | --- | --- | --- | --- |
| 93.15 | 7.00 | $9.34 \times 10^2$ | $8.30 \times 10^2$ | $0.90 \times 10^{-5}$ |
| 113.15 | 5.70 | $9.33 \times 10^2$ | $9.70 \times 10^2$ | $0.63 \times 10^{-5}$ |
| 133.15 | 4.90 | $9.31 \times 10^2$ | $1.1 \times 10^3$ | $0.48 \times 10^{-5}$ |
| 153.15 | 4.20 | $9.31 \times 10^2$ | $1.23 \times 10^3$ | $0.37 \times 10^{-5}$ |
| 173.15 | 3.70 | $9.29 \times 10^2$ | $1.36 \times 10^3$ | $0.29 \times 10^{-5}$ |
| 193.15 | 3.30 | $9.27 \times 10^2$ | $1.5 \times 10^3$ | $0.24 \times 10^{-5}$ |
| 213.15 | 3.00 | $9.25 \times 10^2$ | $1.65 \times 10^3$ | $0.20 \times 10^{-5}$ |
| 223.15 | 2.80 | $9.24 \times 10^2$ | $1.72 \times 10^3$ | $0.18 \times 10^{-5}$ |
| 233.15 | 2.60 | $9.23 \times 10^2$ | $1.80 \times 10^3$ | $0.16 \times 10^{-5}$ |
| 243.15 | 2.50 | $9.22 \times 10^2$ | $1.88 \times 10^3$ | $0.14 \times 10^{-5}$ |
| 253.15 | 2.40 | $9.20 \times 10^2$ | $1.96 \times 10^3$ | $0.13 \times 10^{-5}$ |
| 263.15 | 2.30 | $9.19 \times 10^2$ | $2.03 \times 10^3$ | $0.12 \times 10^{-5}$ |
| 273.14 (ice) | 2.14 | $9.17 \times 10^2$ | $2.11 \times 10^3$ | $0.11 \times 10^{-5}$ |
| 273.16 (water) | 0.56 | $1.00 \times 10^3$ | $4.22 \times 10^3$ | $0.01 \times 10^{-5}$ |
| 283.15 | 0.58 | $1.00 \times 10^3$ | $4.19 \times 10^3$ | $0.01 \times 10^{-5}$ |
| 293.15 | 0.60 | $9.98 \times 10^2$ | $4.18 \times 10^3$ | $0.01 \times 10^{-5}$ |
| 303.15 | 0.62 | $9.96 \times 10^2$ | $4.18 \times 10^3$ | $0.01 \times 10^{-5}$ |
| 313.15 | 0.63 | $9.92 \times 10^2$ | $4.18 \times 10^3$ | $0.02 \times 10^{-5}$ |

**Figure 5:**
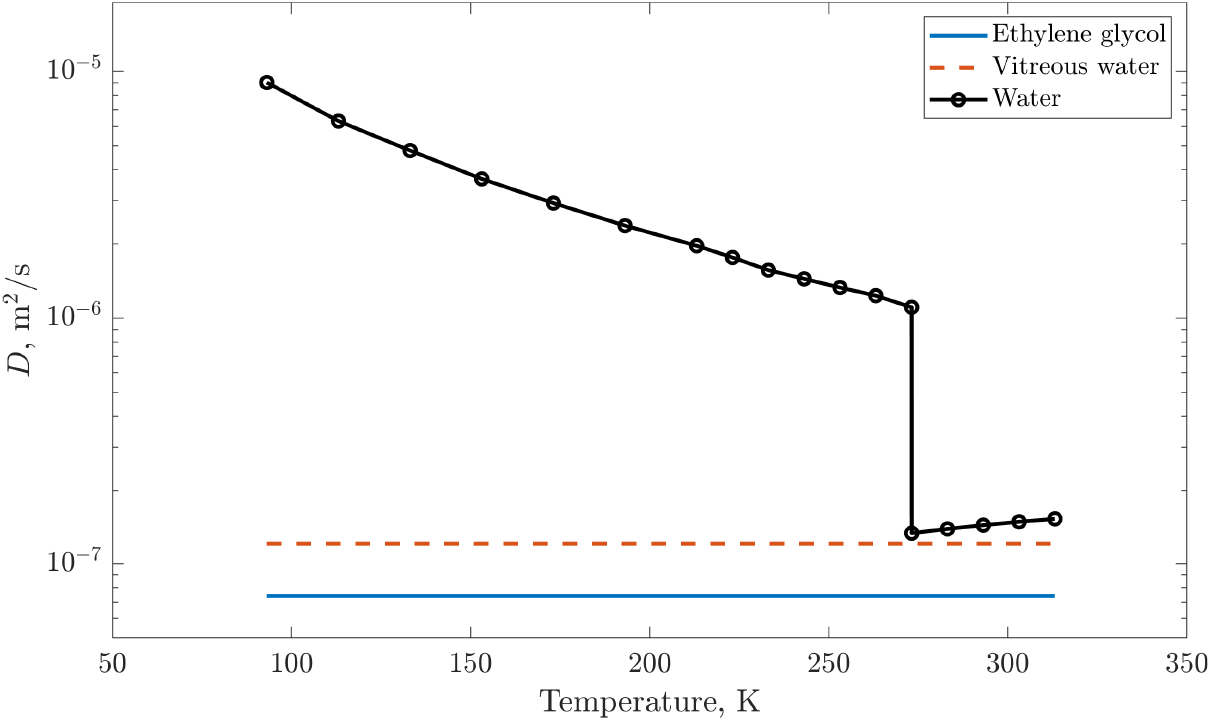
Diffusion coefficient for each of the three different thermophysical parameter choices in Section 2.2.3: ethylene glycol, vitreous water and water (with variable thermophysical parameters). The vertical axis is plotted on a log scale.

Another parameter of interest is the surface heat transfer coefficient, *h*, which along with thermal conductivity, affects the rate of heat transfer at a boundary involving a fluid. The boundary condition arises from Newton’s law of cooling, relating to objects being cooled by forced convection [9], so *h* is dependent on the fluid dynamic behaviour at the interface between the polypropylene plate/cryoprotectant droplet and the liquid nitrogen. The nature of this heat transfer changes depending on whether the liquid nitrogen is in the film boiling regime, or in the nucleate boiling regime. In the film boiling regime, the very large heat gradient between the device and liquid nitrogen at the surface of the model forms an insulating barrier [29], leading to a slower heat transfer rate. Eventually, as the model cools, nucleate boiling will re-establish itself, [32]. In numerical models, these different boiling regimes are often represented by changing the value of *h* [32, 40]. Choosing values of *h* that are determined experimentally can therefore validate the assumption 2.2.1, as we can replicate realistic results without needing to model the underlying fluid mechanics of the system.

In this work, *h* = 2,000 W/m^2^K is chosen, as used in other numerical models of cooling of the Open Cryotop^®^ [31]. This value of *h* matches experimental cooling rates derived using a specialised thermocouple device [25].

A value of interest is the characteristic timescale, which describes how long it takes heat to diffuse throughout a homogeneous material, and is denoted by

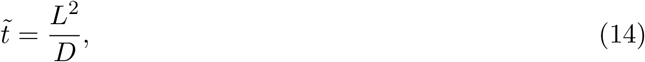

where *L* is the ‘characteristic length’ of a system, *D* is the associated diffusion coefficient and 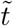 is the timescale. Taking the characteristic length to be the droplet radius, *L* = 0.21mm, and using the *D* value for ethylene glycol in Table 1, a timescale of approximately 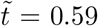 is reported. If 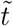 is the dominant timescale in the vitrification process, vitrification times (and therefore cooling rates) will be sensitive to the rate at which the boundary warms, such that cooling rates match those from previous models of the Open Cryotop^®^ [31]. This suggests that the choice of *h*, which determines the rate at which the droplet boundary is cooled, is important in determining the cooling rate of the sample inside the droplet.

Although choices for the thermophysical parameters have been made, this model constitutes a framework, in which any choice of thermophysical parameters can be easily implemented. As more accurate and applicable thermophysical parameter data become available, this model could be quickly rerun to yield more accurate results.

#### 2.2.4. *Cryoprotectant droplets are hemispheres with radius* 0.21 *mm*

Graphics from the user manual of the Open Cryotop^®^, shown in Figure 2, depict one possible shape of the droplets as being concave following the aspiration of excess media. Additionally, the motion of sliding the pipette back may have an effect on the shape of the droplet [16]. In this work, aspiration and pipette motion are ignored, and it is assumed that droplets are spherical caps, as spheres minimise the surface tension in a droplet as a result of the isoperimetric inequality in three dimensions [21]. Observations suggest spherical caps are a good assumption, however, as there is low adhesion of the cryoprotectant to the plate, which means that small volumes relative to high surface tension appear to force the droplet back into a sphere after the pipette is removed.

The contact angle of the droplet must also be described. The contact angle, *θ*, is the angle formed between the plate and the tangent plane to the sphere surface at the point of contact, such that *θ* ∈ (0, π). At this point there is contact between three domains: the fluid droplet, *F*, the solid plate, *P*, and the liquid nitrogen, *Ln*. A tension force *γ_A,B_* exists between each pair of domains, *A* and *B*, and these tension forces may be used to define contact angle through a relation known as Young’s Equation [2],

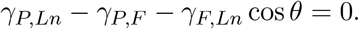

The tension forces depend on the chemical properties of each domain, and environmental factors such as temperature, making it difficult to model them accurately. It has been observed in the lab that the contact angle is at most π/2, as the droplets do not appear truncated at their base [16]. Additionally, the droplet acts structurally as a solid during vitrification [4], so the contact angle does not change as a function of temperature and thus may be taken as a constant. The droplet must have a height greater than 0.2mm above the plate in order to cover the samples, and for a fixed height of droplet, droplet volume decreases with increasing *θ*. A minimal loading volume is desired such that decrease of the cooling rates from increased droplet volume is minimal, so the model takes *θ* = π/2. A radius as 0.21mm is a valid choice, as it may contain a sample and has a volume of 0.019*μ*l, which is within the accepted limit of 0.1*μ*l maximal volume prescribed by the Open Cryotop ^®^ manual [20]. Similar modelling approaches use a droplet whose volume is equal to 0.1*μ*l [33], but choosing a smaller volume allows us to increase the number of samples used without exceeding this upper volume limit.

#### 2.2.5. Samples adopt the chemical properties of the cryoprotectant solution

During vitrification, much of the water volume is removed from the sample by osmosis, falling as low as 5% within two minutes [35], and by artificial collapse [8]. As such, the sample takes absorbs some volume of the surrounding cryoprotectant solution, so the chemical composition of the sample becomes more similar to that of the surrounding solution [12]. This assumption is justified by highlighting that the sample has a very small volume, so unless samples act as strong insulators, or conductors, any change invoked by this assumption is minimal. This assumption is made indirectly in previous literature, in which only a droplet without a suspended sample was modelled [31, 33].

### 2.3. Spatial arrangements

When constructing spatial arrangements of samples, the samples are placed such that they just touch each other, or there exists only a very small separation (see Figure 6), so that the effect of tightly packing samples together can be tested. Placing samples as close to each other as possible also maximises the overlap between the separate droplets formed around each sample, which ensures that the extra cryoprotectant needed to cover both samples is minimal, reflecting the vitrification guidelines recommending a minimum volume of cryoprotectant [20].

**Figure 6:**
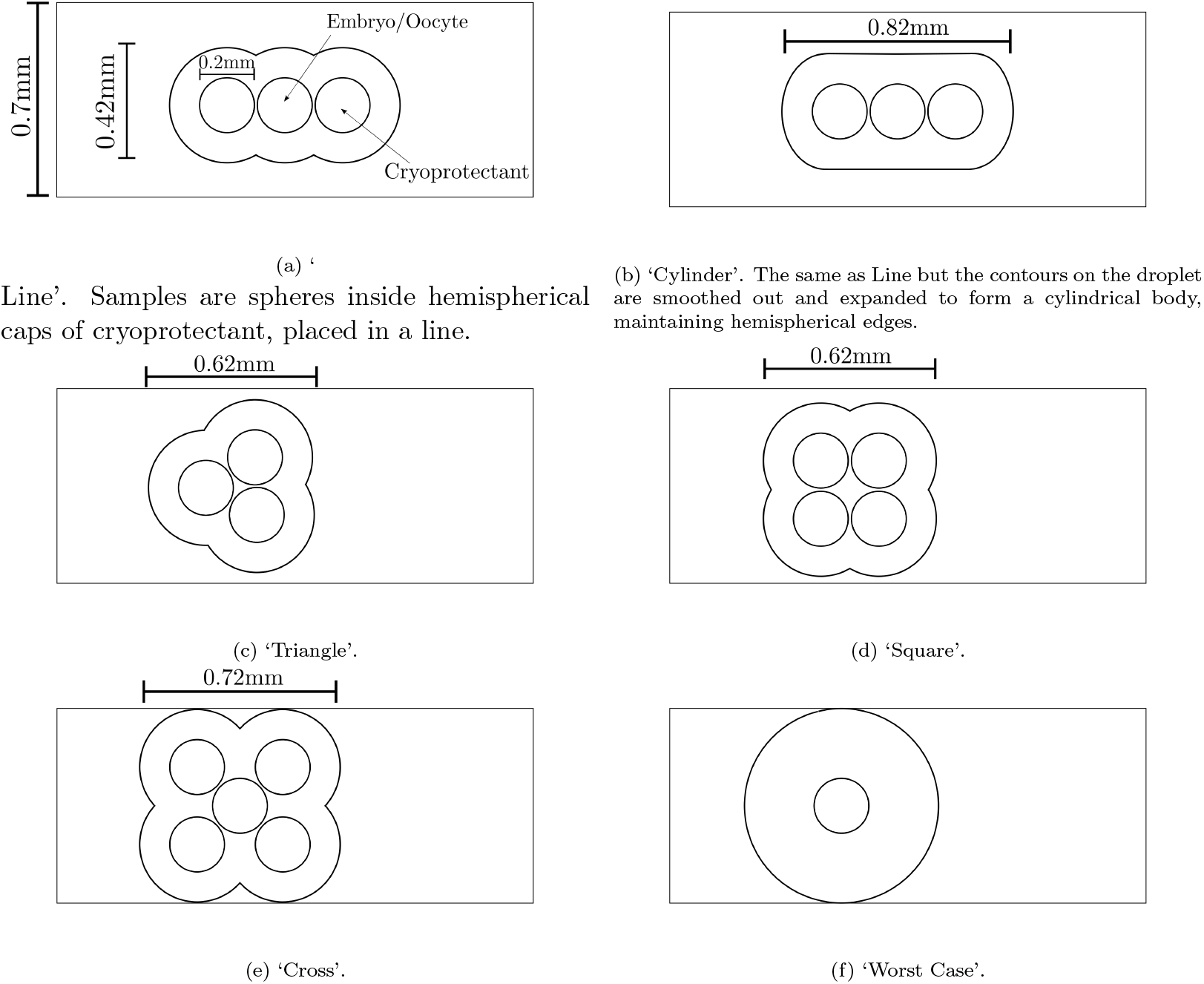
Top-down visualisations of cases explored by our model.

The first case studied in this work is a single sample, so that we provide a reference case for more complicated geometries. Up to three more samples are then added in a line running down the length of the plate, to study the effect of increasing the number of samples (see Figure 6a).

In the next case studied, instead of forming a cryoprotectant droplet from the union of the droplets around it, the transfer of multiple samples leads to a small increase in the cumulative volume of droplets merging, and hence samples are assumed to be in a ‘Cylinder’ of cryoprotectant with rounded ends (Figure 6b). This case is considered in order to test how minor perturbations in droplet geometry compared to the ‘Line’ case might alter cooling rates.

Additionally, other cases are studied where the samples are not aligned along the centre of the plate, but instead congregate into more complex geometries, which is sometimes observed when multiple samples are vitrified. Three different arrangements are considered, a ‘Triangle’ (Figure 6c), a ‘Square’ (Figure 6d) and a ‘Cross’ (Figure 6e). In practice the actual locations of the samples on the plate may vary, but cases for every possible location of the samples cannot be constructed. As such, only some of the most representative densely packed cases possible are examined.

In the Triangle case, samples are placed such that their centres lie on the corners of an equilateral triangle with side length 0.2mm. In the computational model, such an arrangement may cause issues where COMSOL Multiphysics 5.5 cannot efficiently generate a mesh around the contact points, which appear to overlap. As such, in practice the leftmost sample is shifted slightly to prevent mesh generation failures (see Figure 6c). In the Square case, sample centres form the corners of a square whose side lengths are 0.2mm. In the Cross case, an arrangement similar to that of the square is desired, but with a sample in the centre. This arrangement would cause the droplet to hang off the plate, however, so samples are instead placed on the corners of a rectangle with side lengths 0.28mm wide by 0.3mm long, leaving enough space to place a sample in the centre of the rectangle. The measurements shown in Figure 6 are calculated based off of these rules.

Manufacturer’s guidelines allow a maximum of four samples per device [20], which means that the Cross case is not considered a valid arrangement within an IVF clinic. This case is tested nonetheless because it is an extreme case that allows us to quantify the effect on the rate of cooling on a sample that is surrounded by other samples, so that the validity of the guidelines may be confirmed.

The sixth case considered is the ‘Worst Case’ benchmark, in which the droplet has radius equal to the width of the plate (Figure 6f), and contains only a single sample at its centre. The volume of this droplet is 0.09*μ*l, which is just less than the recommended maximal volume of 0.1*μ*l, and as such, this represents a case which is known to be safe as it fits within current guidance [20]. As the droplet volume is maximised, this will represent the case with the slowest cooling rate still considered to be viable for sample vitrification. Whilst the model cannot necessarily be used to predict a minimum safe, or effective, cooling rate, any case in which cooling rates are greater than those observed in the Worst Case simulations can be considered to have sufficiently fast vitrification.

### 2.4. Computational modelling

A Finite Element Method (FEM) is used to simulate the temperature throughout the models, solving Equations (4)–(13). Employing COMSOL Multiphysics 5.5, adaptive meshes are constructed for each given geometry to balance accuracy and computational cost. The computational model discretises the domain into individual elements, the temperature of which can be defined and calculated at each time point of interest. A mixture of triangle and tetrahedral elements are used, as determined by the solver. A ‘fine’ element discretisation is employed, which uses a minimum of 2182 tetrahedral elements and 1320 triangular elements. The temperature is calculated over 2 seconds at 0.02 second intervals, and the simulation run time is only a few seconds.

The average temperature within the samples is calculated as a volume average in COMSOL Multiphysics 5.5, by integrating the temperature within the samples, and dividing by the sample volume [11]. Such volume integrals can be expressed in terms of the domain volume *V* as [9]

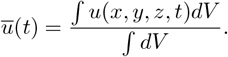

The code used in this work is available on Github, https://github.com/OstlerT/MultipleEmbryoModels.

## 3. Results

Figure 7 shows the average temperature over time of the samples in the cases in which between one and three samples are positioned running along the centre of the plate (cases visualised in Figure 6a and Figure 6a), with separate plots for each set of assumed thermophysical parameters. Figure 8 shows the temperatures in the Triangle, Square, Cross and Worst Case arrangements, with a minimum of three samples included. The time taken to reach a temperature of 143.15K, for all cases, is shown in Table 3. The largest time to vitrification is 0.45s, the associated cooling rates are displayed in Table 4.

**Figure 7:**
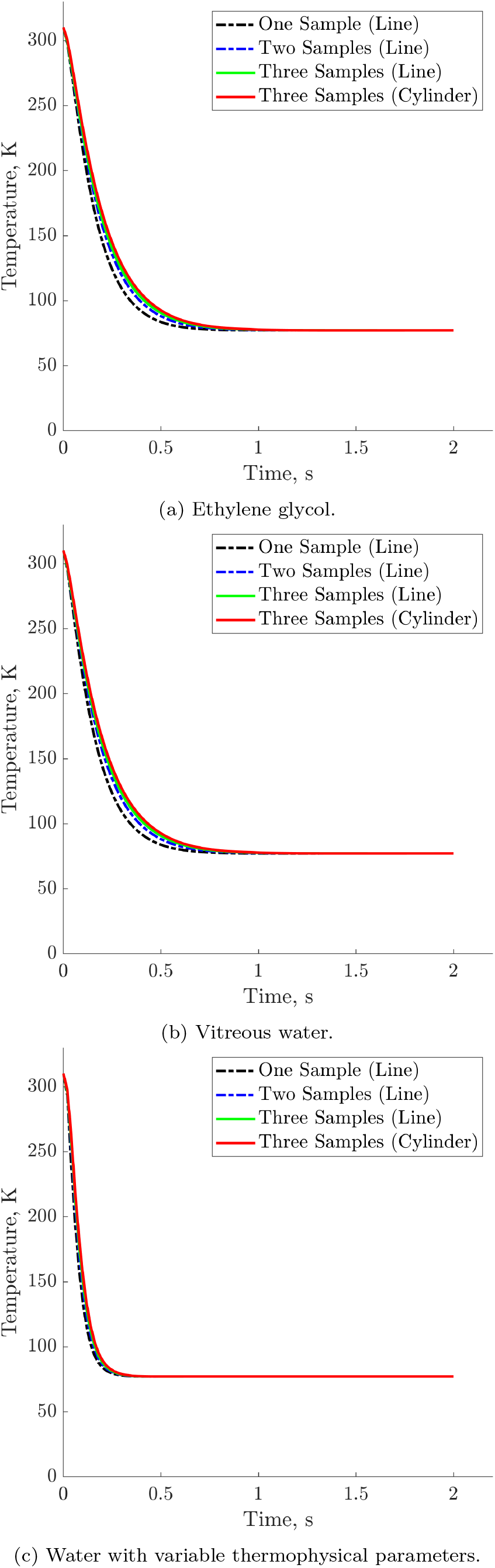
Average temperature over time. The number of samples increases in a linear format (Figure 6a), with the bounding cylindrical case also presented (Figure 6b).

**Figure 8:**
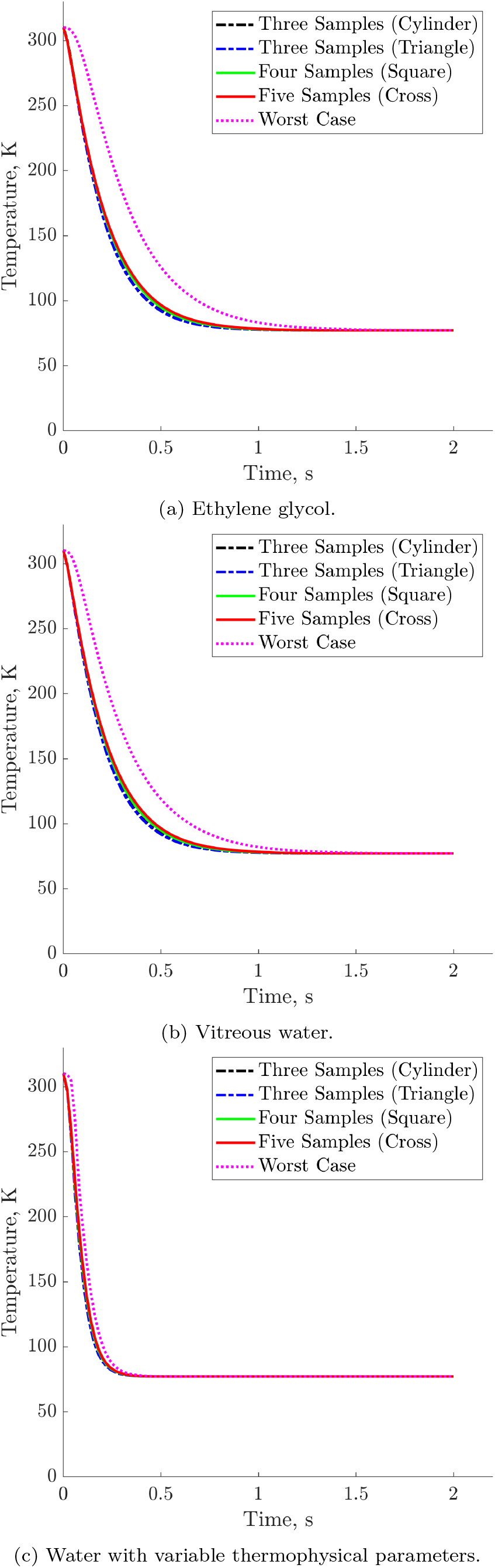
Average temperature over time for different sample arrangement patterns, depicted in figures 6c-6f.

**Table 3:** Time taken in seconds for average temperature to reach 143.15K inside the samples for all choices of thermophysical parameters, either ethylene glycol (EG), vitreous water (vitW) or water with variable thermophysical parameters (W).

| Model | Number of samples | Time, s |  |  |
| --- | --- | --- | --- | --- |
|  |  | EG | vitW | W |
| 1 Line | 1 | 0.26 | 0.26 | 0.14 |
| 2 Line | 2 | 0.28 | 0.28 | 0.16 |
| 3 Line | 3 | 0.30 | 0.30 | 0.16 |
| Cylinder | 3 | 0.30 | 0.30 | 0.16 |
| Triangle | 3 | 0.30 | 0.30 | 0.16 |
| Square | 4 | 0.32 | 0.32 | 0.16 |
| Cross | 5 | 0.32 | 0.32 | 0.16 |
| Worst Case | 1 | 0.48 | 0.44 | 0.18 |

**Table 4:** Cooling rates from 310.15K to 143.15K in the simulations, using cooling times from Table 3. Results shown for all choices of thermophysical parameters, either ethylene glycol (EG), vitreous water (vitW) or water with variable thermophysical parameters (W).

| Model | Cooling rate, K/min |  |  |
| --- | --- | --- | --- |
|  | EG | vitW | W |
| 1 Line | 38541 | 38541 | 71576 |
| 2 Line | 35788 | 35788 | 62629 |
| 3 Line | 33402 | 33402 | 62629 |
| Cylinder | 33402 | 33402 | 62629 |
| Triangle | 33402 | 33402 | 62629 |
| Square | 31314 | 31314 | 62629 |
| Cross | 31314 | 31314 | 62629 |
| Worst Case | 20876 | 22774 | 55670 |

The results in Figure 7 demonstrate that increasing the number of samples leads to a decrease in the cooling rate. The temperature is similar in all cases, and increasing the number of samples in a line will not affect the cooling rate of a sample in the centre of the line, as samples now lose heat through their contact surfaces with the liquid nitrogen much faster than they would lose heat to the adjacent samples. Hence, placing more than three samples arranged in a line would not be of value, to the 3 Line case is therefore representative of any greater number of samples placed in a line.

Of particular note in Figures 7a and 7b is the minor difference between the two cases with three samples, 3 Line and Cylinder. The additional cryoprotectant in the Cylinder case slows down cooling of the samples by a small amount, but from Table 3 it can be seen that this difference is on the order of magnitude of milliseconds. Therefore, it can be concluded that these two cases are almost identical, which supports assumption 2.2.4, that minor perturbation in droplet shape is unimportant.

Figure 8 shows the temperature over time in cases where arrangements are not all aligned along the centre of the plate. A similar effect as in Figure 7 is observed; increasing the number of samples leads to a corresponding decrease in the cooling rate. In the two cases with three samples, the Cylinder case and the Triangle case, the profiles are almost identical, whilst adding more samples to the model increases the time to vitrification.

The average temperature in the Worst Case in Figure 8 displays the smallest cooling rate out of all cases considered. Given that this particular construction is designed to be the most accurate ‘worst-case’ scenario that is still considered safe to undertake, it can be concluded that any arrangement whose cooling rate is larger than that of Worst Case is also a safe and suitable case. As such, all of cases tested may be considered safe to use with regards to cooling rates.

In all cases using the temperature-dependent thermophysical parameters of water, shown in Figures 7c and 8c, we find that the effect of varying the sample number and arrangements is smaller than when using constant parameters. We have already demonstrated in Figure 5 that for variable thermophysical parameters, the diffusion coefficients are significantly higher at low temperature, which implies that the samples will cool much faster. This is despite the increasing thermal conductivity, which suggests that the rate at which the boundary warms up should decrease (Equation (3)). The simulations in Figures 7c and 8 demonstrate that even when considering variable thermophysical parameters, the sample arrangements still yield similar spatio-temporal thermal profiles; the conclusion is confirmed in this particular case.

The cooling rates in Table 4 are of similar order of magnitude to those reported by Kitazato^®^ at 23, 000K/min. Despite using the same thermophysical parameters for vitreous water as in [31], we report smaller cooling rates. The reason for this discrepancy is as follows. The droplets used in [31] are also spherical caps, but with a smaller contact angle, so they have a height of 0.1mm and a width of 0.35mm. This yields a droplet volume of approximately 0.02*μ*l, which means that the Line case with one sample presented above, with a droplet volume of also approximately 0.02*μ*l, should give the closest comparison, yet has approximately half the cooling rate. This can be attributed to assumptions regarding the geometry of the model. In [31], a 2D axis symmetric geometry is assumed, which means the plate volume is smaller and has a greater surface area to volume ratio, increasing the rate at which the plate cools relative to the model we use here. This effect is demonstrated when tracking the warmest point in each model; in this work, the warmest point is always at the contact between the sample and the plate, whilst in [31] the warmest point is raised above the plate, implying heat transfer through the plate is more significant than in our model. Additionally, the plate material is assumed to be polyethylene [31], with a diffusion coefficient (thermal diffusivity) of 1.4 × 10^−7^m^2^/s, which is a little under twice that of polypropylene, as assumed in this work (see Table 1), which further contributes to increased cooling through the plate. Additionally, the droplets used in [31] have a smaller contact angle, and as such have approximately 1.5 times the surface area. With a greater surface area, the droplets cool much faster for the same volume.

The difference between the results of our model and experimental cooling rates [25] can be attributed to the use of constant thermophysical parameters, which as previously discussed, yield overestimations in the time taken for cooling. When assuming variable thermophysical parameter, cooling rates are much higher (4). Though these cooling rates are likely to be excessively high, given the cryoprotectant does not consist of pure water, the model determines upper and lower cooling rates for the constant and variable thermophysical parameter simulations, respectively, both of which are within the correct order of magnitude for experimental results for this device.

## 4. Discussion

The problem of measuring cooling rates during vitrification of multiple oocytes or embryos is an important one, with direct clinical implications. As it cannot be addressed through experiments, in this modelling work we have shown that differences in cooling rates induced by different arrangements of samples are not large enough to affect survival rates of samples undergoing vitrification, so long as droplet volumes remain constant and within manufacturer’s limits. All arrangements we have simulated have higher cooling rates than would be observed in the laboratory, represented by our Worst Case simulations; therefore they are considered to be valid configurations. The small perturbations in cooling times observed between cases would be within the error tolerance of experimental readings and, thus, although these results may not be quantitatively exact, they do offer a robust “rule of thumb” time scale of around half a second for time to vitrification. This means that there is no need for embryologists to spend time arranging loaded samples on the plate; instead they should focus on aspirating the medium efficiently. This can save precious time for embryologists, and justifies operating to their personal preference within the confines of standard operating procedures.

The differences in the cooling rate observed between our model other models can be attributed to variations in the geometry and assumptions made in the model construction. Despite the small disparity of around 9.2% between our Worst Case cooling rates and those reported by Kitazato [23], our model yields suitable lower and upper bounds for the cooling rates of samples vitrified in varying numbers and spatial arrangements, both of which are within the safe and appropriate bounds reported for the Open Cryotop device. If any spatial arrangement of samples has a larger cooling rate than our extreme Worst Case lower bound, then it may be considered to have a sufficiently high cooling rate for safe vitrification.

Although cooling rates have been used as a comparable measure between this work and pre-existing work in the field, we highlight some issues associated with this measure. First, cooling rates have a non-linear dependence on the predetermined temperature interval used, so where different authors use different temperature intervals, cooling rates cease to be comparable. Additionally, temperature evolution over the system in question occurs within the order of half a second, which means that very small absolute differences in the time taken for cooling can result in extreme variation in the cooling rate, which is an inverse of the time taken for cooling. As such, vastly differing cooling rates may appear to justify selecting one technique or arrangement over another, when in reality vitrification occurs in an almost identical time frame. As such, we recommend caution when relying solely on cooling rates as a definitive measure of the quality of vitrification.

The results of this work are dependent on the validity of the assumptions made. The assumption in Section 2.2.4 is that all droplets must be exact hemispheres, but this may not be the case in reality. Different-shaped droplets such as those depicted in [33], or ovoid shapes, may exist. In this case, the arrangement of samples may create different-shaped droplets in different cases, which would lead to different characteristic lengths and therefore different cooling rates. Such perturbations to the simulations would lead to small differences in our predicted results.

The question of how many samples should be mounted to a single device is becoming increasingly important, and this study shows that the only limitation from a thermal perspective is the skill of the embryologist. Specifically, they may mount any number of samples that can be properly stored within 0.1*μ*l of vitrification solution within a reasonable time frame, with suitable in-house validation. Our work does not assess other factors associated with loading of multiple samples, such as the time taken to process samples, or cost efficiency, it only validates that loading of multiple samples per device is thermally justifiable. As a result, it cannot be concluded that vitrification of multiple samples, or samples from any given arrangement is optimal. Instead, it can be concluded that thermal differences are an unlikely explanation for variable survival rates arising from variable sample number and arrangement.

Further work can be done to refine the model and achieve greater accuracy. This work focuses only on the Open Cryotop^®^ device, but could be replicated for other techniques and devices in common usage. The easiest way to refine the model is to revisit the assumptions in Section 2.2. Specifically, the assumptions 2.2.2 and 2.2.4, about the size and shape of the samples and droplets, can be easily altered. These geometries could be further refined to be more accurate and account for realistic droplet shape, larger droplet volumes, and variable sample location and size within the droplets.

One might also wish to refine the assumption in Section 2.2.1, which suggests liquid nitrogen is a stationary isothermal liquid. In practice, the liquid is boiling, which results in additional mechanisms of temperature change, such as boiling-induced convection and other boiling specific effects [29]. The effect of these additional mechanisms between may differ between cases and may affect cooling rates. This would involve more complex numerical modelling which would account for fluid motion and phase change.

Further work could also be undertaken to explore the choice of parameters in the model. Parameter choices are taken from literature and COMSOL Multiphysics 5.5, but may not reflect accurate parameter values. Additionally, it was assumed that samples have the thermal properties of their surrounding medium. The choice of the surface heat transfer coefficient is of interest. Making a good choice of this parameter is highly important to making the model as accurate as possible. In our model, it is trivial to replace the values of the parameters, so as more accurate parameter values become available for the materials and compounds associated with the problem at hand, the model can be updated to generate more accurate approximations of cooling rates.

Another important factor to be considered is that some models suggest that maximising cooling rates is not the optimum approach to improving survival, [7] and that there is instead an optimal range for cooling rates balanced by the risk of damage to cells from the cryoprotectant for high cooling rates and the risk of intracellular ice formation for low cooling rates. As such it may not be appropriate to look for arrangements that yield high cooling rates, but instead arrangements that give cooling rates within some optimal range. Additionally, there is evidence that cooling rates are less significant for the survival of oocytes than warming rates [25]. Indeed, it may be that some balance between cooling rates and warming rates should be sought. As such, important future work could be to also model the effect of spatial arrangement on the warming process.

Despite the highlighted shortcomings of our work, the simulations provide the correct order of magnitude, in line with experimental findings. Specifically, as long as the amount of cryoprotectant does not exceed the maximum volume given by the manufacturers’ guidelines, the cooling process should be on the order of half a second, or less, regardless of the spatial arrangement or number of mounted samples, which means that all arrangements considered in this work exhibit high enough cooling rates to facilitate successful vitrification.

## 5. Acknowledgements

This research is funded by the Knowledge Economy Skills Scholarship (KESS2). KESS2 is a pan-Wales higher level skills initiative led by Bangor University on behalf of the HE sector in Wales. It is part funded by the Welsh Government’s European Social Fund (ESF) convergence programme for West Wales and the Valleys.

The authors also thank the reviewers of this paper, for their in-depth analysis and careful, constructive criticism, which has significantly improved the manuscript.

